# A Physiological Characterization in Controlled Bioreactors Reveals a Novel Survival Strategy for *Debaryomyces hansenii* at High Salinity and Confirms its Halophilic Behavior

**DOI:** 10.1101/2020.01.10.901843

**Authors:** Clara Navarrete, August T. Frost, Laura Ramos-Moreno, Mette R. Krum, José L. Martínez

## Abstract

*Debaryomyces hansenii* is traditionally described as a halotolerant non-conventional yeast, being the model organism for the study of osmo- and salt tolerance mechanisms in eukaryotic systems for the past 30 years.

However, unravelling of *D. hansenii’s* biotechnological potential has always been difficult due to the persistent limitations in the availability of efficient molecular tools described for this yeast. Additionally, there is a lack of consensus and contradictory information along the recent years that limits a comprehensive understanding of its central carbon metabolism, mainly due to a lack of physiological studies in controlled and monitored environments. Moreover, there is controversy about the diversity in the culture conditions (media composition, temperature and pH among others) used by different groups, which makes it complicated when trying to get significant conclusions and behavioural patterns.

In this work, we present for the first time a physiological characterization of *D. hansenii* in batch cultivations using highly instrumented and controlled lab-scale bioreactors. Our findings contribute to a more complete picture of the central carbon metabolism and the external pH influence on the yeast ability to tolerate high Na^+^ and K^+^ concentrations. Finally, the controversial halophilic/halotolerant character of this yeast is further clarified.

## Introduction

*Debaryomyces hansenii* is described as a halophilic/halotolerant non-conventional yeast. It is globally present in seawater and has been isolated from soil, air, materials of plant and animal origin as well as from polar waters and ice from Antarctic and Artic glaciers (Norkrans 1966; Breuer and Harms 2006; Gunde-Cimerman *et al.* 2009). *D. hansenii* has been a model for the study of osmo- and salt tolerance mechanisms in eukaryotic cells over the last 30 years (Adler *et al.* 1985; Prista *et al.* 1997, 2005). Potassium and sodium are crucial factors for yeast growth in high salt environments, and its halotolerant nature has been fully confirmed by the fact that the presence of sodium in the medium protects the yeast cells against oxidative stress and additional abiotic stresses like extreme pH or high temperature (Almagro *et al.* 2000; Papouskova and Sychrova 2007; Navarrete *et al.* 2009).

*D. hansenii*’s genome was completely sequenced in 2004 (Dujon *et al.* 2004), and although more than 6500 genes have been annotated since then, the molecular characterization of this yeast is still far from being fully known. The majority of these annotated genes are mainly associated with salt and osmotic stress tolerance mechanisms (Prista *et al.* 2016). Besides, it has been described that *D. hansenii* has the highest coding capacity among yeasts (79.2% of the genome) with 6906 detected coding sequences (CDS) and a gene redundancy of almost 50% (Gènolevures online database, igenolevures.org/databases/).

As mentioned before, potassium and sodium fluxes (and their accumulation in cell organelles) play a key role in ion homeostasis and halotolerance in *D. hansenii*. Genes for K^+^ influx (*DhTRK1*, *DhHAK1*) were identified and studied at a molecular, transcriptional and protein level by different authors (Prista *et al.* 2007; Martinez *et al.* 2011). Two different plasma membrane cation efflux systems (DhEna1/2, DhNha1) have also been described (Almagro *et al.* 2001; Velkova and Sychrova 2006) as well as two intracellular Na^+^/H^+^ antiporters (DhKha1 and DhNhx1) (Garcia-Salcedo *et al.* 2007; Montiel and Ramos 2007). Very importantly, the role of glycerol production and accumulation in the osmotic stress response to high osmolarity was also fully established by Gustafsson and Norkrans in 1976, and further confirmed by Adler *et al.* in 1985.

From a biotechnological point of view, *D. hansenii* is considered of great interest, due to its salt tolerant character (Gadanho *et al.* 2003; Butinar *et al.* 2005; Prista *et al.* 2005; Ramos *et al.* 2017 among others), especially in food industry where it is used in the ripening process of sausages by production of exopeptidases, development of flavor characteristics, and production of cheeses among others (Lopez Del Castillo-Lozano *et al.* 2007; Cano-Garcia *et al.* 2014). Moreover, *D. hansenii* is known to respire a broad range of carbon substrates and produce mycocins against other yeast species, like *Candida* (Banjara *et al.* 2016).

Nevertheless, only a few genes with high biotechnological relevance have been characterized so far in *D. hansenii*. It is the case of *DhJEN1*, coding for a monocarboxylic acid transporter (Casal *et al.* 2008), and genes coding for xylitol reductase, xylose dehydrogenase and xylose/H^+^ transporter (Biswas *et al.* 2013; Ferreira *et al.* 2013), all of them used for the production of second-generation bioethanol in the fermentation of pentoses by *Saccharomyces cerevisiae* (Breuer and Harms 2006).

The difficulties when studying *D. hansenii*’s biotechnological potential have always been related to the limitations in the availability of highly efficient molecular tools described for this yeast (Prista *et al.* 2016). There is also a lack of information and full understanding of its carbon metabolism and physiological characterization during biotechnological processes.

On top of that, the large variation in culture conditions used (e.g. media composition, temperature, and pH) in previous studies impedes thorough understanding and solid conclusions on this peculiar yeast’s behaviour. To date, nobody has conducted an in-depth physiological analysis of this yeast in a controlled environment (e.g. using bioreactors), and there is a lack of consensus about its capacity to produce ethanol in high saline environments or its cell performance in the presence of high salt concentrations. While some studies point to a beneficial role of salts on *D. hansenii’s* performance (Almagro *et al.* 2000; Papouskova *et al.* 2007; Navarrete *et al.* 2009; Garcia-Neto *et al.* 2017), others claim that sodium is detrimental in terms for the fitness of this yeast in general (Capusoni *et al.* 2019; Sánchez *et al.* 2018). In addition, the osmo- and halotolerant terminology are often used incorrectly, as there are some studies in which sodium is used for both purposes, despite there are reported evidences about sodium and potassium triggering a different effect. Potassium does not cause a toxicity response, whereas sodium has been described to have detrimental effects on cell growth at lower concentrations than potassium, pointing to some authors to recently claim *D. hansenii* not being halophilic but just halotolerant (Sánchez *et al.* 2018), whilst several previous studies report *D. hansenii* as an halophilic yeast (Gonzalez-Hernandez *et al.* 2004; Chao *et al.* 2009; Martinez *et al.* 2011).

In relation to *D. hansenii*’s alcoholic fermentation capacity, the kinetics of cell inactivation in the presence of ethanol at 20%, 22.5% and 25% (v/v), have been measured by progressive sampling and viable counting, and used as an inference of the ethanol resistance status of different yeasts (Pina *et al.* 2004). *D. hansenii* PYCC 2968T (a.k.a. CBS 767, which we use in our present study) was found to be one of the most sensitive yeasts to ethanol in this study. Ethanol (or other polyols) production has been only reported when a mix of sugars or complex substrates are used as carbon source. For example, *D. hansenii* CCMI 941 was cultivated in xylose/galactose or xylose/glucose (Tavares *et al.* 2008) for production of xylitol, although only ethanol and glycerol were produced for a xylose/glucose ratio above 30%. Another strain background, *D. hansenii* B-2, was grown in semi-synthetic banana peel-yeast extract-peptone broth for ethanol production, where 40% (w/v) of glucose was fermented to 5.8% of ethanol (Brooks, 2008). In 2009, Calahorra *et al.* stated the activation of glucose fermentation by salts in *D. hansenii* Y7426. In this work, the authors incubated yeast cells in media containing 40mM of glucose, and afterwards they made a protein extraction and measured the ethanol production *in vitro*, by the enzyme extract. Only marginal ethanol production was reported though, within the range of micromoles per gram of glucose. However a most recent work performed by Garcia-Neto *et al.* in 2017, indicates otherwise: that the presence of high salts increases *D. hansenii’*s respiratory activity, in the same strain (Y7426). Overall, no ethanol production by *D. hansenii* has been shown during the growth process using glucose as a sole carbon source.

In this work, we present for the first time a physiological characterization of *D. hansenii* during batch cultivations in highly instrumented and controlled bioreactors. *D. hansenii*’s carbon metabolism and the external pH influence on the yeast capacity to tolerate high Na^+^ and K^+^ concentrations are also shown. Finally, its capability of ethanol production and the controversial halophilic/halotolerant character of this yeast are further discussed, and a novel survival strategy at high saline environments suggested.

## Materials and Methods

### Strain and culture conditions

The *Debaryomyces hansenii* strain CBS767 (PYCC2968; Prista *et al.* 1997; Navarrete *et al.* 2009) was used in this study. For the micro-fermentation experiments and growth assay on plate, IBT22 and IBT26 (CBS1101) strains were also used (from IBT culture collection of fungi from DTU, Denmark). Glycerol stocks containing sterile 30% glycerol (Sigma-Aldrich, Germany) were used to maintain the strain, and were preserved at −80⁰C.

Yeast extract Peptone Dextrose (YPD) medium plates with 2% agar, were used for growing the cells from the cryostocks at 28°C. For the pre-cultures of yeast cells, synthetic complete medium was used (6.7 g/L Yeast Nitrogen Base w/o amino acids, from Difco, plus 0.79 g/L complete supplement mixture, from Formedium). Separately sterilized 2% D-(+)-glucose monohydrated (VWR Chemicals, Germany) was added to the medium, and pH was adjusted to 6.0 with NaOH. All the solutions were autoclaved at 121°C for 20 min. Cells were incubated in 500 mL baffled Erlenmeyer shake flasks (culture volume 100 mL) at 28°C, 150 rpm for at least 24 hours.

For the growth curves in low glucose conditions, *D. hansenii* pre-cultures were prepared as specified in the above paragraph. From those pre-cultures, cells were grown in the same medium but with 0.2% of glucose, and with or without 1M/2M of NaCl or KCl, in order to get their growth profile. Initial OD_600_ in the flask was 0.1 and samples were taken during eight days of cultivation at 28°C, 150 rpm.

### Growth assay on plate

Three *D. hansenii* strains (CBS767, IBT22 and IBT26) were pre-cultured for at least 24 h in synthetic complete medium (pH 6) at 28°C as specified in “Strain and culture conditions” section, and cell suspensions were prepared from those pre-cultures. Cell drops at OD_600_ 0.1 and serial dilutions (1:10, 1:100) were spotted on YNB plates (pH 6) with or without KCl or NaCl at different concentrations (0.5, 1 and 2M). Cell growth was monitored at 28°C for 144 h or 240 h (when higher salt concentrations were used). Pictures were taken using an in-house build digital camera set-up (Nikon D90 SLR camera).

### Microscopy

*D. hansenii* cells were grown on synthetic complete medium (pH 6) with or without KCl or NaCl (0.5, 1 and 2M) at 28°C as specified in “Strain and culture conditions” section. Undiluted cell samples were taken and photographed at the microscope (BA310E & camera 580INT, In Vitro DK) after 51 h of cultivation.

### Bioreactor cultivations

Batch cultivations were performed in biological replicates (between 2-7 per condition) in 1.0 L Biostat Qplus bioreactors (Sartorius Stedim Biotech, Germany). The temperature was controlled at 28°C and pH was maintained at 6.0 (when desired) by the automatic addition of 2M NaOH / 2M H_2_SO_4_, and measured by pH sensors (Model EasyFerm Plus K8 160, Hamilton). The volumetric flow rate (aeration) was set at 1 vvm and the stirring was constant at 600 rpm. Dissolved oxygen concentration was also measured by DO sensors (Model OxyFerm FDA 160, Hamilton). The working volume in the vessel was 0.5 L, using exactly the same medium composition as in the pre-culture, containing either 2% or 0.2% of glucose when required. To study the effect of salt in *D. hansenii* cells, NaCl or KCl (PanReac Applichem, ITW Reagents) were added to the medium before autoclavation. The bioreactors were inoculated with 24 h inoculum from the pre-culture to get an initial OD_600_ of 0.05-0.1. Samples for dry weight, optical density and HPLC were taken after the CO_2_ percentage values reached 0.1 and until stationary phase.

### Micro-fermentation experiments

All micro-fermentations were carried out in BioLector^®^ I and II systems (m2p Labs, Germany) using 48-well FlowerPlate^®^ (MTP-48-B, m2p Labs, Germany) with a working volume of 1.5 mL. Synthetic complete medium (Yeast nitrogen base 6.7 g/L, Complete Supplement Mixture 0.79 g/L, D-(+)-Glucose monohydrated 2%) at different pH (4, 6 and 8) was used, and different temperatures were tested (26, 28 and 30°C). The pH was adjusted with NaOH, and potassium phthalate monobasic ≥99.5% (20.4 g/L, Sigma-Aldrich, Germany) was added to the media and used as a pH buffer for the experiments. Three *D. hansenii* strains (CBS767, IBT22 and IBT26) were pre-cultured in YNB medium (pH 6) at 28°C for at least 24 h, washed with ddH_2_O and inoculated in the corresponding wells at an initial OD_600_ 0.1. The programmed settings were as follows: filter; biomass (gain 10 and 3, respectively for the two BioLector^®^ systems), humidity; on (>85% using ddH_2_O), temperatures; 26, 28, and 30 °C, oxygen supply; 20.85% (atmospheric air), agitation speed; 1000 rpm. The biomass formation was monitored for at least 70 h.

The light scattering data obtained from the BioLector^®^ I and II systems were analyzed using R version 3.6.1 (The R Foundation for Statistical Computing).

### Metabolite analysis

The concentrations of glucose, glycerol and ethanol were measured by High Performance Liquid Chromatography (Model 1100-1200 Series HPLC System, Agilent Technologies, Germany). The injection volume was 20 μl, the eluent 5mM H_2_SO_4_ and the flow rate was set at 0.6 mL/h. The temperature of a Bio-Rad Aminex HPX-87H column was kept at 60°C. A standard solution containing glucose (20 g/L), glycerol (2 g/L) and ethanol (20 g/L) was used (Sigma-Aldrich, Germany) for exo-metabolites concentration determination.

### Off-gas and dissolved oxygen measurements

CO_2_ and O_2_ concentrations were continuously analyzed in real time by mass spectrometry coupled to the off-gas line (model Prima PRO Process MS, Thermo Scientific, UK). From the off-gas CO_2_ emission data, given in percentage, the maximum specific growth rate was calculated. Off-gas CO_2_ and O_2_ emission data were also used to determine the Carbon Dioxide Evolution Rate (CER), the Oxygen Uptake Rate (OUR) and the Respiratory Quotient (RQ).

Dissolved oxygen values were measured by DO sensors (Model OxyFerm FDA 160, Hamilton) as previously described in “Bioreactor cultivations” section.

### Analytical procedures

Specific growth rates in the different growth conditions were calculated based on the optical density (OD_600_) and emitted CO_2_ values.

Yield coefficients and carbon balances were used to describe the metabolite, by-products and biomass formation by the yeast cells, and were calculated based on the DW, accumulated CO_2_ and HPLC data. The average minimal formula CH_1.79_O_0.50_N_0.20_ for yeast dry cell biomass composition was used for the calculations as proposed by Roels in 1983. The specific glucose consumption rates were calculated based on the logarithmic method proposed in Görgens *et al.* in 2005.

### Statistical analysis

Statistical analysis was performed with Microsoft Excel® 2016 (version 1903, 32-bit, USA). All values are represented as averages ± 95% confidence interval of independent biological replicate cultures. For regression analysis, the coefficient of determination (r^2^) was used to determine the statistical significance of the fit, where a value above 95% was considered statistically significant.

## Results

### *D. hansenii* growth rate under the effect of high salt concentrations

Specific growth rates were calculated based on the optical density and the volumetric CO_2_-production rates under the different growth conditions tested (Fig. 1). In general terms, a higher maximum specific growth rate was observed when a higher salt concentration is present in the media, except for concentrations of 2M of either salt, compared to control conditions. Sodium exhibit a significantly stronger positive impact than potassium, as the highest growth rate was reached in the presence of 0.5-1M NaCl (similar observations were made for KCl, where concentrations of 0.5M gave slightly higher values than 1M, both resulting in a higher growth rate compared to control, although lower than for NaCl). On the other hand, a decrease in specific growth rate is observed when the concentration of salts is above 1M, and up to 2M (Fig. 1, tables). However, even at concentrations of both sodium and potassium up to 1.5M, the growth rate values are still significantly higher than in control conditions (no salts added).

**Figure 1.**
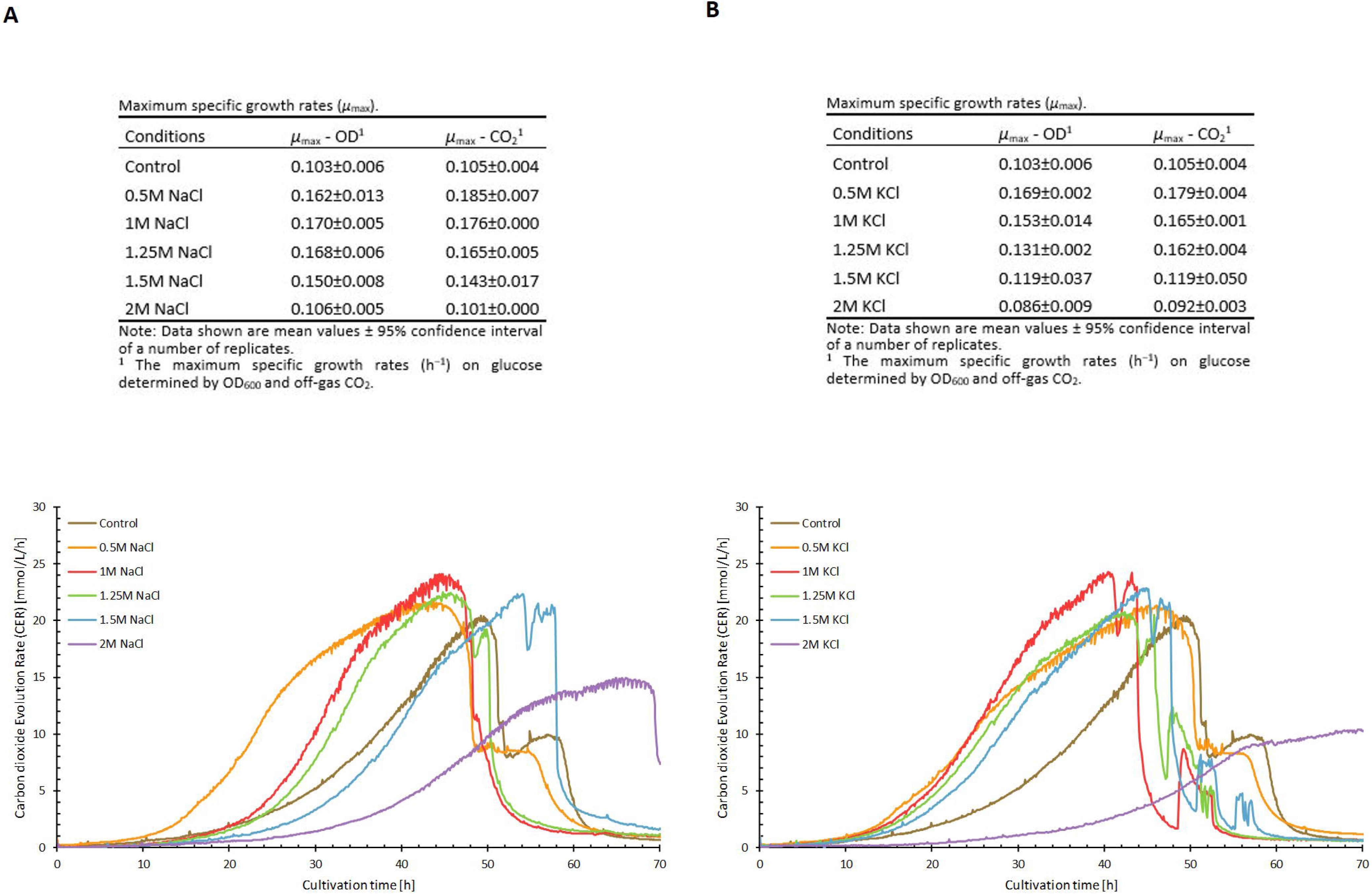
Maximum specific growth rates and carbon dioxide evolution rate (CER) profiles from batch cultivations with sodium (**A**) or potassium (**B**) chloride. Maximum specific growth rates determined by optical density (OD) and off-gas CO_2_ emission data from batch cultivations of *Debaryomyces hansenii* in synthetic complete media at 28°C and pH 6 with varying concentrations of sodium/potassium chloride are shown in the tables. The CER profiles are based on off-gas CO_2_ emission data over time from one replicate for each condition and are representative of their associated replicate(s).

Surprisingly, at 2M NaCl the growth rate is lower, but still very close to control conditions, although a prolonged lag phase is observed in comparison (exponential growth phase starts around 30-35 hours in the control vs. 40-45 hours for 2M NaCl, as inferred from the CO_2_ profiles in Fig.1). In contrast, the addition of 2M KCl results in a lower μmax compared to the 2M NaCl and also lower than the control, reinforcing the fact that NaCl still exerts some beneficial effect in *D. hansenii*’s cell performance overall. Microscope samples were taken, randomly, from the liquid cultures at increasing salt concentrations and also from control cultures (no salts added), and neither morphological changes nor in cell size were appreciated among all the conditions (Suppl. Fig. S1).

The same conclusion can be inferred if we observe the carbon dioxide evolution rate (CER) of *D. hansenii* in the presence of salts over time (Fig. 1). The profiles show a higher CO_2_ production (that can be translated into higher glucose consumption, hence a higher metabolic rate) when the cells grow in the presence of NaCl or KCl compared to control conditions with no salt. Once again, this effect begins to decrease when the salt concentrations are over 1.25M, nevertheless still higher than compared to the control conditions, except for 2M in which the production rate is lower than the control for both salts (Fig. 1).

### Biomass yield on substrate and biomass titers show a differential effect among K^+^ and Na^+^

The observed decrease in the specific growth rate from above 1M of salt seems to be compensated by a slightly higher biomass yield upon glucose, as was observed for the dry weight measurements during the cultivation time, although only for the addition of potassium (Table 1). The final biomass titers were not significantly different though, with the exception of the 2M sodium which were slightly higher in comparison (Suppl. Fig. S2).

**Table 1.** Yield coefficients and specific glucose consumption rates from batch cultivations.

| Conditions | $Y_{SC}^1$ | $Y_{SX}^1$ | $Y_{SE}^1$ | $Y_{SG}^1$ | C-balance<sup>2</sup> | SGCR ($r_s$)<sup>3</sup> |
| --- | --- | --- | --- | --- | --- | --- |
| Control | 0.63±0.01 | 0.35±0.00 | ns<sup>4</sup> | ns | 0.98±0.01 | 0.120±0.004 |
| 0.5M NaCl | 0.57±0.03 | 0.32±0.02 | ns | ns | 0.90±0.05 | 0.157±0.001 |
| 1M NaCl | 0.63±0.07 | 0.35±0.04 | ns | ns | 0.98±0.11 | 0.163±0.012 |
| 1.25M NaCl | 0.60±0.02 | 0.34±0.01 | ns | ns | 0.94±0.04 | 0.159±0.002 |
| 1.5M NaCl | 0.63±0.10 | 0.35±0.06 | ns | ns | 0.98±0.16 | 0.133±0.014 |
| 2M NaCl | 0.63 | 0.36 | ns | ns | 0.99 | 0.098±0.014 |
| 0.5M KCl | 0.61±0.00 | 0.34±0.00 | ns | ns | 0.95±0.00 | 0.153±0.009 |
| 1M KCl | 0.68±0.05 | 0.38±0.03 | ns | ns | 1.05±0.08 | 0.159±0.012 |
| 1.25M KCl | 0.63±0.05 | 0.35±0.03 | ns | ns | 0.98±0.08 | 0.148±0.004 |
| 1.5M KCl | 0.68±0.04 | 0.38±0.02 | ns | ns | 1.06±0.06 | 0.144 |
| 2M KCl | 0.66±0.14 | 0.37±0.08 | ns | ns | 1.02±0.21 | 0.138±0.001 |

Yield coefficients from batch cultivations of *Debaryomyces hansenii* at different salt concentrations are shown in the table. Additionally, the specific glucose consumption rates during the exponential phase calculated based on the logarithmic method are presented. The cells were grown in synthetic complete medium with or without salt at different concentrations, at 28°C and pH 6. Data shown are mean values ± 95% confidence interval of a number of replicates.
<sup>1</sup> Yield of CO<sub>2</sub> (C), Biomass (X), Ethanol (E), and Glycerol (G) from Glucose (S) in cmol/cmol.
<sup>2</sup> Carbon balance on a C-mol basis, where a value of 1 indicate a closed carbon balance.
<sup>3</sup> Specific Glucose consumption rate in cmol/cmol/h based on logarithmic method.
<sup>4</sup> ns: Not statistically significant. The amount of ethanol and glycerol was neglectable.

When the specific glucose consumption rate was calculated, *D. hansenii* showed higher rates of consumption in the presence of NaCl or KCl than in control conditions, being 1M the optimal concentration showing the highest values, and again this effect started to revert once the concentration of salts used was above 1M up to 1.5M, although still higher values than those observed in the control (Table 1), while at 2M NaCl the specific consumption rates were lower than the control. The specific glucose consumption values observed were higher for sodium than for potassium at lower concentrations (0.5M - 1M - 1.25M), as seen before for the specific growth rate, however the opposite is observed at higher concentrations (uptake rates are higher in the presence of potassium).

To further investigate the effect of salts on the specific glucose uptake rates, *D. hansenii* was grown in low glucose conditions (0.2%), preliminarily in shake flasks and later by using bioreactors, to further confirm the findings in flasks. In the shake flask experiments, it was observed that cells growing in the presence of 1M KCl adapted much faster to the nutrient limitation and were able to grow at a significantly higher rate compared to the other conditions tested (Suppl. Fig. S3). This effect was also observed in the presence of 1M NaCl, but at a lower level. Moreover, the presence of 2M of either salt in the medium had no detrimental effect for the cells, which still grew at a similar growth rate compared to the control. Strikingly, the presence of 2M of NaCl resulted in almost 3 times higher biomass concentration than the control conditions. In this particular circumstance (high salts and low glucose), the effect of potassium seems to be beneficial in terms of growth rate, but sodium seems to have a better effect in cell performance in the long run, since the growth rate is slower but the final biomass concentration is much higher in return (Suppl. Fig. S3). All together seem to confirm, without a doubt, the halophilic character of *D. hansenii* in one hand, and the differential effect of Na^+^ and K^+^ salts, in the other.

Based on these results in shake-flasks, and in order to obtain a more complete dataset that would support such observations, we then conducted the same type of batch cultures with a reduced initial glucose concentration (0.2%) in bioreactors without pH regulation (thus mimicking the previous flask conditions). Our previous observations in flasks cultivations were confirmed, however interesting differences were observed. The carbon dioxide evolution rates (CER) exhibit a much faster adaptation (Fig. 2A), evidenced by a shorter lag phase, when cells grow in the presence of 1M of either salt (NaCl or KCl), which confirms our observations in flasks. Further analyses showed significantly higher specific glucose uptake rates and, surprisingly, much higher growth rates at 1M of NaCl or KCl, than compared with the rest of the conditions (Fig. 2B). In contrast, CO_2_ yields and biomass yields are higher at concentrations of 2M, being 2M NaCl the highest values of all conditions compared. Additional data showing the timeline of DW, OD_600_ and glucose consumption, are shown in Suppl. Fig. S4.

**Figure 2.**
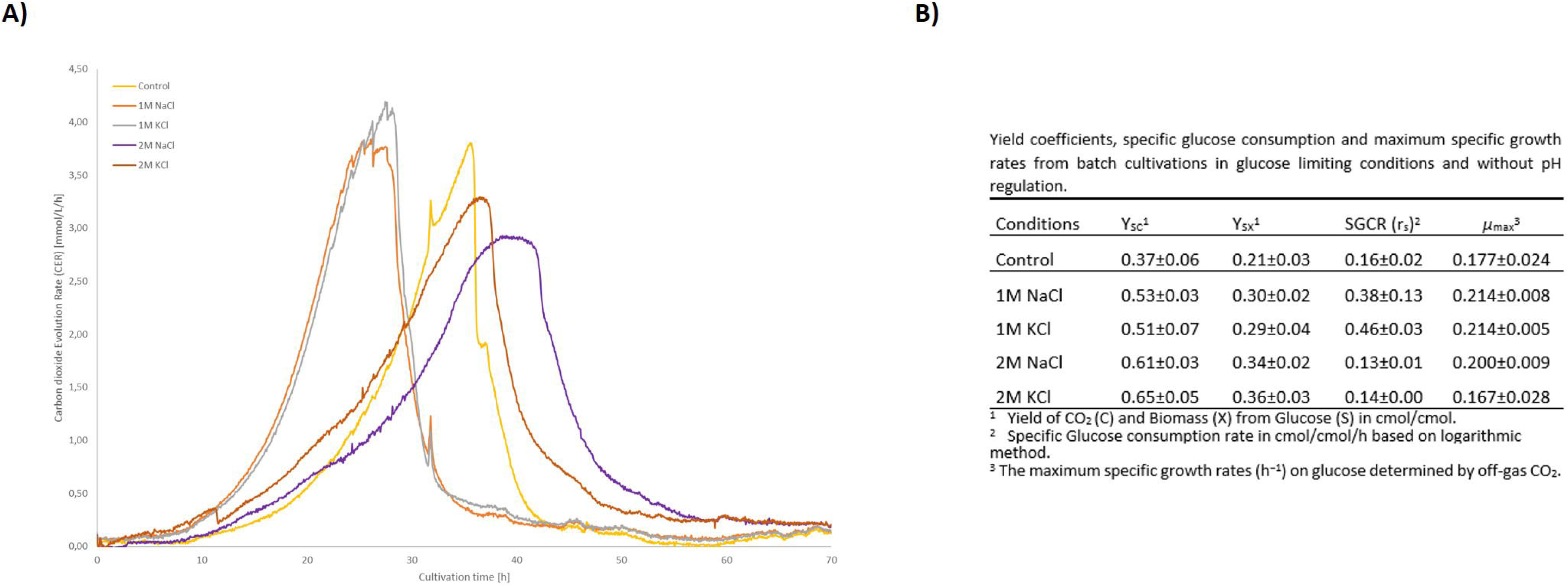
Carbon dioxide evolution rate (CER) profiles and yield coefficients, specific glucose consumption and maximum specific growth rates from batch cultivations under limiting glucose conditions and without pH regulation. Off-gas CO_2_ emission data from batch cultivations of *Debaryomyces hansenii* in synthetic complete media containing 0.2% of glucose at 28°C and initial pH value of 6 with varying concentrations of sodium/potassium chloride are shown in the figure (**A**). The CER profiles are based on off-gas CO_2_ emission data over time from one replicate for each condition and are representative of their associated duplicate. Yield coefficients and maximum specific growth rates from batch cultivations of *Debaryomyces hansenii* at different salt concentrations are shown in the table. Additionally, the specific glucose consumption rates during the exponential phase calculated based on the logarithmic method are presented (**B**). The cells were grown in synthetic complete medium containing 0.2% of glucose and with or without salt at different concentrations, at 28°C and initial pH of 6. Data shown are mean values ± 95% confidence interval of duplicates.

As observed in the graph, the CER profiles are very similar between control conditions and cells growing at 2M KCl, only for 2M NaCl the CO_2_ production peak seems lower (Fig. 2A), however when we look at the CO_2_ yields on glucose, they are higher for both salts at 2M in comparison to 1M and also with the control. The biomass yields at 2M are also higher than the 1M concentration, and much higher than the control, as shown in Figure 2B. Altogether, this confirms our observations in the shake flasks, pointing that on reduced glucose conditions, the positive effect of the presence of high salt concentration is even more accused. Moderate levels of salt (1 M) result in higher glucose consumption rates, and higher specific growth rates, while higher concentration of salts (2M) result in higher biomass yields and CO_2_ yields, at the cost of a lower growth rate. A metabolism switch that can be described as: from growing as fast as possible to growing as much as possible. We also further confirmed that sodium exerts a more positive impact than potassium, once again.

It is worth mentioning that the observed final OD values were higher for the control in bioreactor experiments than in shake flasks. Still we observed a higher biomass when cells are growing in 2M NaCl, as we also saw in the shake flasks, however cell growth in control conditions was arrested at a lower OD in flasks compared to bioreactors, so the difference in total biomass is less significant for the latter (Suppl. Fig. S3). This is not surprising though, and simply illustrates the importance of using well stirred reactors in physiology studies.

### Dissolved oxygen concentration and RQ levels confirm a fully respiratory metabolism, discarding a fermentative process in *D. hansenii* in our experimental conditions

Dissolved oxygen levels during bioreactor cultivations were observed to decrease faster when Na^+^ or K^+^ is present in the medium, being those levels lower while the concentration of salt increases, suggesting a higher oxygen demand (Fig. 3). This confirms a higher metabolic activity at increasing salt concentrations, which again points to the halophilic character of *D. hansenii*, and might already suggest a fully respiratory metabolism, regardless of the presence or absence of (high) salt in the cultivation media. It is worth mentioning that in some of the tested conditions the dissolved oxygen levels gets very closed to 0, however this does not mean that cells were suffering from oxygen limitation: the airflow inlet throughout the whole bioreactor cultivation was constant, so the cells were never deprived from oxygen, as can be seen (Fig. 3) in the very stable RQ value represented by a constant flat line (RQ = 1) which coincides with the period of higher oxygen utilization. Thus, the lower dissolved oxygen values simply illustrate a much higher oxygen utilization by the cells, but never oxygen deprivation (which would show then an RQ value > 1, and it is not our case).

**Figure 3.**
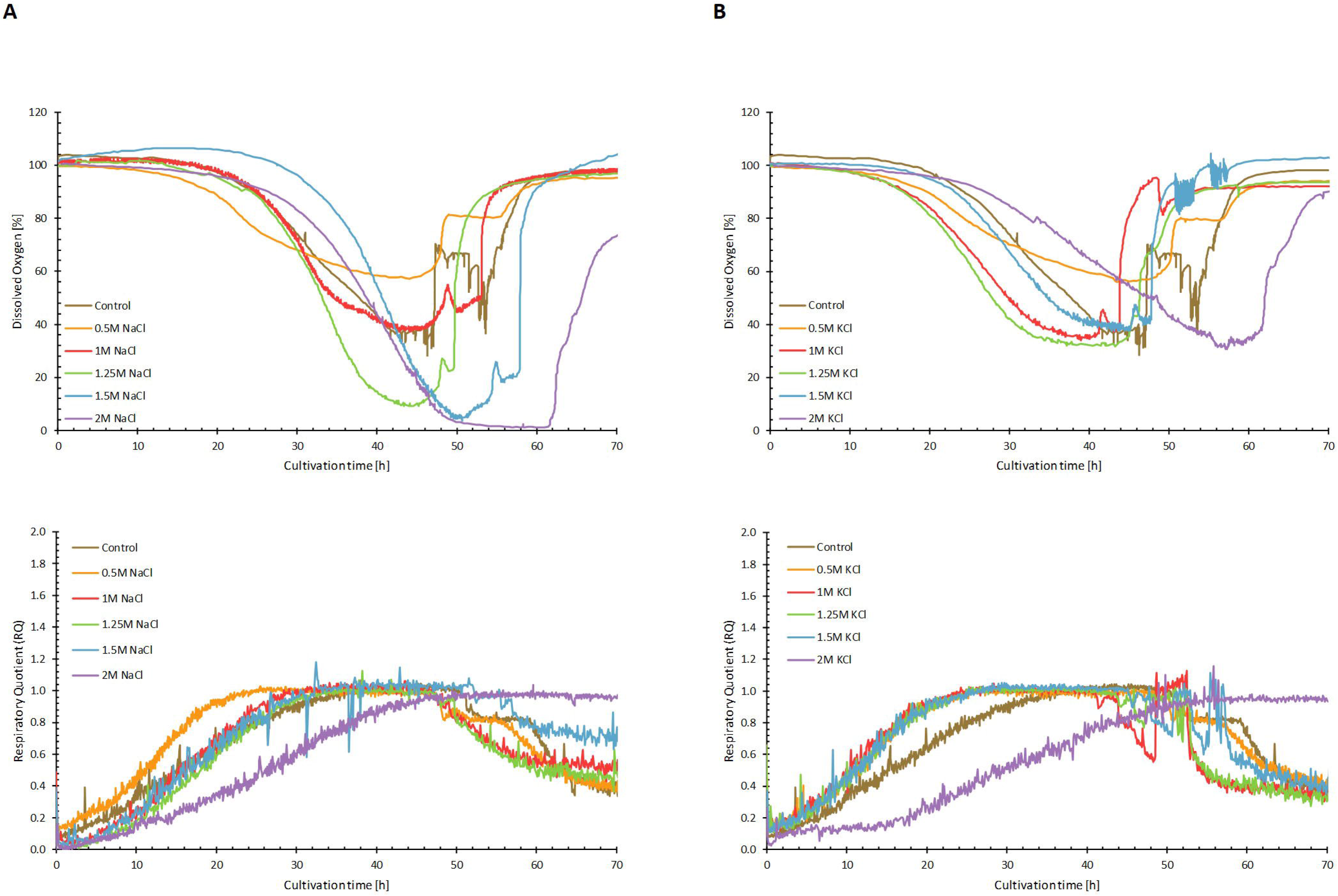
Dissolved oxygen (DO) and respiratory quotient (RQ) profiles from batch cultivations with sodium (**A**) or potassium (**B**) chloride. Dissolved oxygen (%) levels measured over time (h) in synthetic complete media at 28°C and pH 6 with varying concentrations of sodium/potassium chloride, are represented in the figure. The DO profiles are based on one replicate for each condition and are representative of their associated replicate(s). The RQ profiles are based on OUR and CER values calculated from off-gas O_2_ consumption and off-gas CO_2_ emission data from one replicate for each condition and are representative of their associated replicate(s).

The Respiratory Quotient (RQ) values calculated over the entire course of the culture, show to be below or equal to 1 during the exponential growth phase, but never above this value (Fig. 3), this further supports the absence of fermentation, and therefore confirms that *D. hansenii* is not producing ethanol from glucose in our conditions, not even in the presence of high salts, as reported by Calahorra *et al.* (2009). As a final confirmation of the absence of a fermentative process, our off-gas data and the HPLC analysis show no trace of ethanol neither in the gas phase nor in the liquid broth samples (Table 1).

Although it worths mentioning that this previous work (Calahorra *et al.* 2009) was performed using a different *D. hansenii* strain in other culture conditions, and their measurements correspond to ethanol production using enzyme extracts from previously cultured cells, a follow up study performed by García-Neto *et al.* (2017) contradicted such observation using the same strain. Our observations align well with the latter study, confirming that no ethanol production is occurring in our strain either, thus further proving that *D. hansenii* is crabtree negative.

### Additional limiting factors affect cell performance in non-controlled cultivation environments

In order to test the influence of external pH regulation in the halotolerant / halophilic behavior of *D. hansenii*, parallel bioreactor cultivations in normal glucose (2%) were run, in which no pH control was set, and extracellular pH levels were measured on-line in real time. It was observed that, when no pH control was exerted, the CO_2_ profiles of *D. hansenii* evidenced a long maintained plateau phase which cannot be seen in pH-controlled fermentations, both in control conditions and under the effect of salts (Fig. 4). The changes in the external pH over time, for the different conditions tested, are also shown in Supplementary Fig. S5.

**Figure 4.**
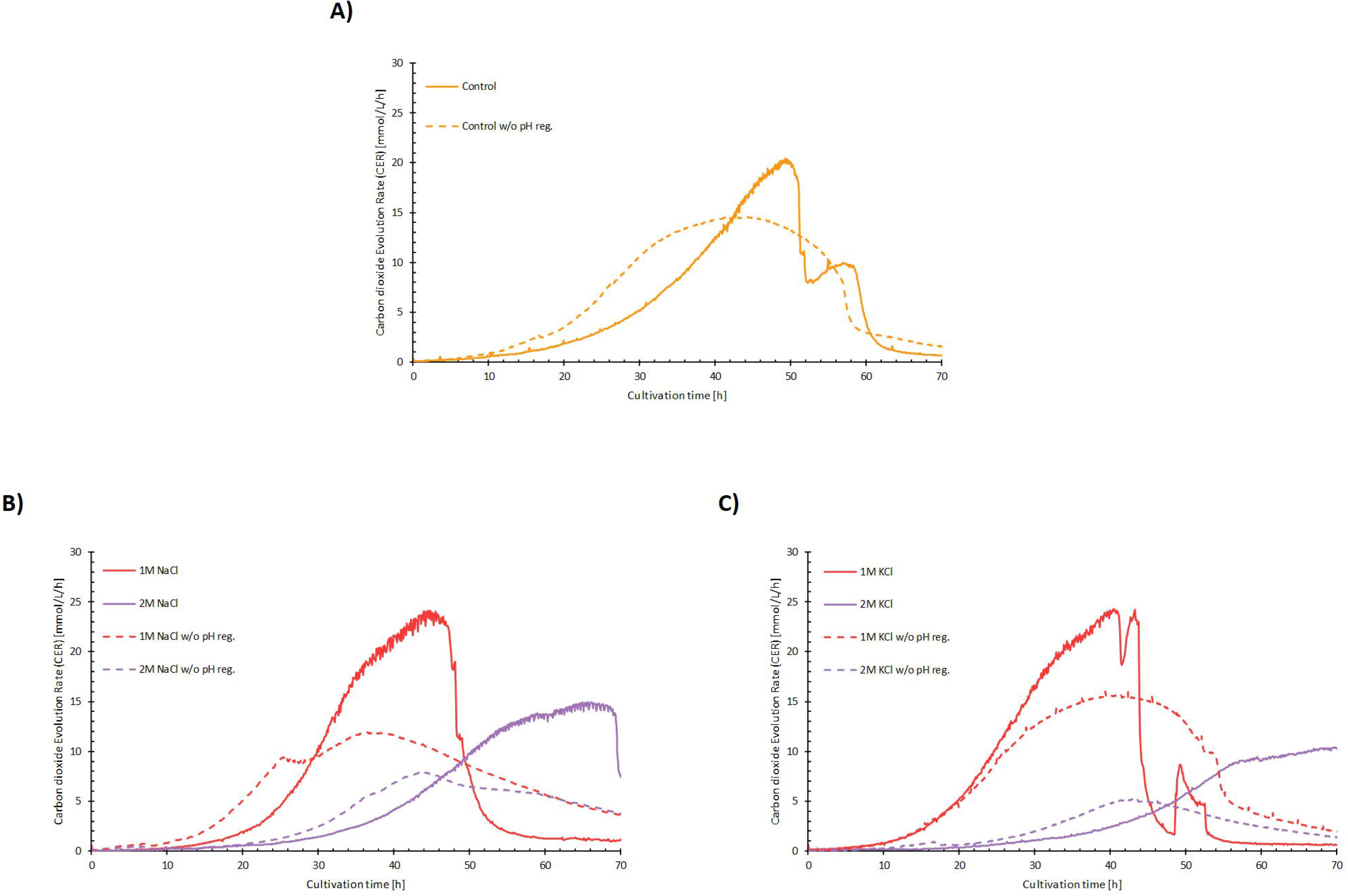
Graphical comparison of CER profiles with and without pH regulation for batch cultivations with varying concentrations of salts. The CER profiles are based on off-gas CO_2_ emission data over time from one replicate for each condition and are representative of their associated replicate(s). The cells were grown in synthetic complete media at 28°C with and without pH regulation (pH 6 with regulation). The cells were either grown without salts (**A**), NaCl (**B**) or KCl (**C**).

This led us to determine, that no meaningful conclusion about metabolic patterns or behaviour can be made, with a high degree of accuracy, in non-controlled cultivation environments: whatever conclusion made is undoubtedly linked to other limiting factors. This means, that previous studies reporting such conclusions, whose data was obtained from non-controlled environments (such as shake flasks, for example) must be considered cautiously.

If we have a look at the maximum specific growth rates under no pH regulation, it can be observed an increase in the μ_max_ when 1M NaCl and KCl are used. This points to a potential summative effect of low pH and high salt concentrations and, once again, sodium seems to have a higher positive impact on growth when compared to potassium. This had been already suggested by Almagro *et al* (2000), so our results further confirm this observation. The specific growth rates decrease once the concentration of both salts is close to 2 M, as also observed in pH-controlled experiments, although this time there is a lower growth rate compared to the control conditions, than when pH control is set in the bioreactors (Table S1 at Supp. material).

Interestingly, the previously described plateau in the CO_2_ profiles is not observed for non-pH controlled bioreactors run with glucose limiting conditions, where sharper and well defined peaks can be seen in the CO_2_ emission profiles (Figure 2A). Here, a faster pH decrease occurs in optimal growth conditions, corresponding to 1M salt added (either NaCl or KCl) to the media (Suppl. Fig. S6), compared to the control. This further supports the summative effect of low pH and increasing salts, to be very beneficial for overall *D. hansenii’*s performance, as already shown in Figure 2B.

About the type of limitation that is occurring, and that generates the long-plateau phase seen in the CO_2_ emission profiles at normal glucose (2%), and why we do not observe such limitation in limited sugar (0.2%) we cannot elaborate further with the data that we have. This shall need to be addressed by future, and more specific, experimental studies.

### Spot-tests and micro-fermentation screening analyses confirm the advantages of high salinity, especially sodium, for the optimal proliferation of *D. hansenii’s* strains

In a follow up round of experiments, we aimed to confirm our observations in liquid media for all the conditions tested (including controlled bioreactors and shake flask cultivations), that is: a faster growth at medium-high salinity (1M salts) evidenced by higher growth rates, and slower rate but longer proliferation capacity to higher biomass densities at higher salinity (up to 2M). To that end, we performed spot tests in solid media plates with increasing salt concentrations (Figure 5), this time including other *D. hansenii* strains than the model CBS767 to check if our observations are strain dependent or otherwise if our conclusions can be extrapolated to other strains of the same species, corresponding to other isolates from other international collections.

**Figure 5.**
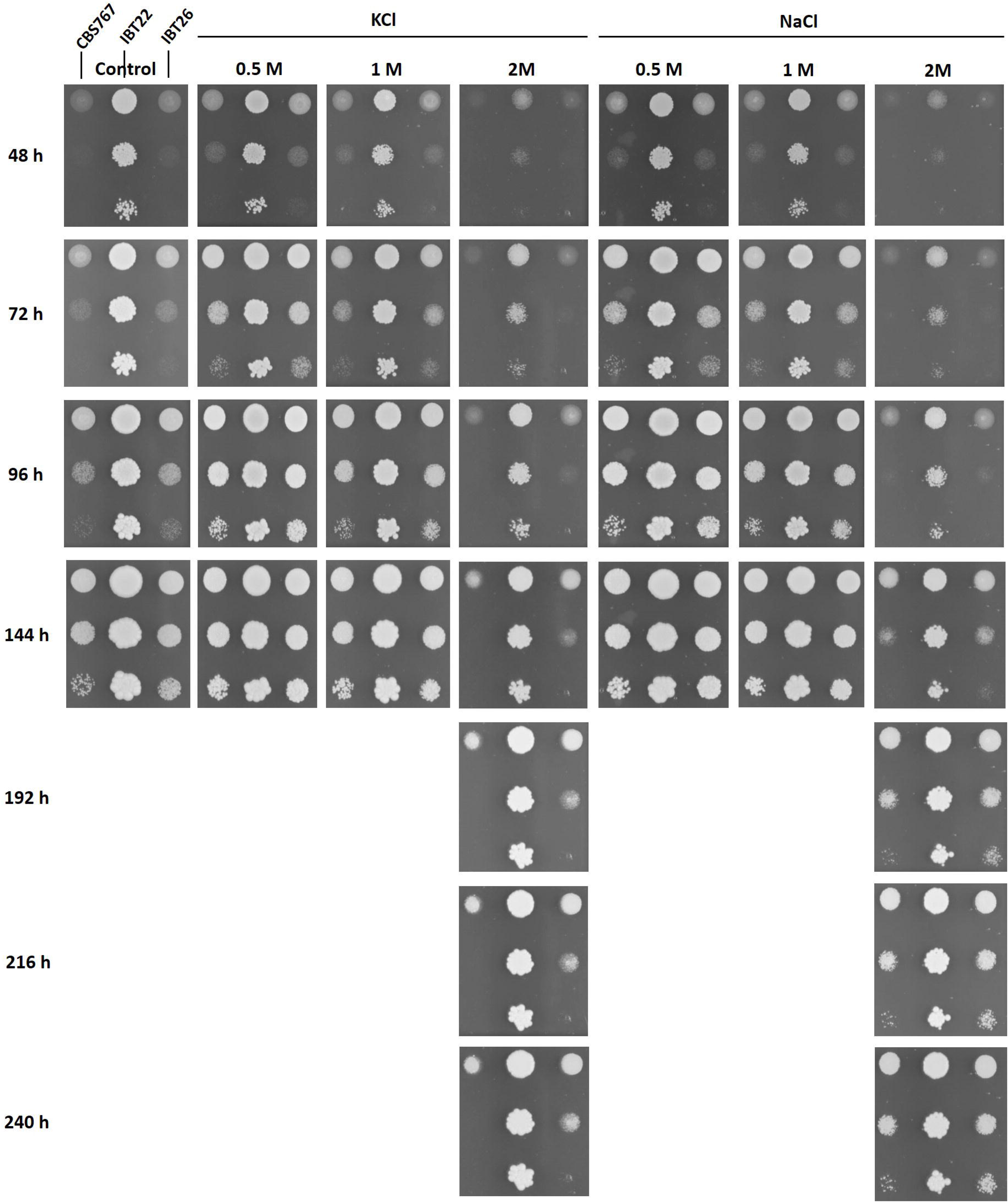
Growth assay of *D. hansenii* strains CBS767, IBT22 and IBT26 in YNB plates (pH 6) control and supplemented with different concentrations of KCl or NaCl. Cells were grown in flasks without salt for 24 h, and cell suspensions were prepared from those cultures. Cell drops at OD_600_ 0.1 and serial dilutions (1:10, 1:100) were spotted on the plates. Growth was monitored for 144 h, or 240 h when higher salt concentrations were used.

As shown in the figure, we confirmed in one hand that the presence of NaCl results more beneficial than KCl as observed in the previous experiments in liquid media, and on the other hand that moderate-to-high concentrations (0.5 – 1M) of sodium make yeast grow much faster than the control, whilst high concentrations (2M) result in slower growth. However in the long run (240 hours of cultivation), we can observe that the yeast proliferation capacity remains unaltered, reaching similar cell densities than the control. This further confirms that sodium is definitely not detrimental at all as it has no inhibitory effect on *Debaryomyces*’ proliferation capacity.

Finally, complementary tests were performed in controlled micro-fermentors (BioLector^®^) including the other *D. hansenii* strains, to assess the effect of both salts in combination with other abiotic parameters such as different pH conditions (4-6-8) and temperatures (26-28-30 °C). The use of such micro-fermentation systems allows both to have controlled conditions and at the same time screening several parameters in parallel reactions, thus studying the influence of such combinations and offering the possibility of identification of behavioural trends among the yeast strains tested, by using surface-response plots (Figure 6).

**Figure 6.**
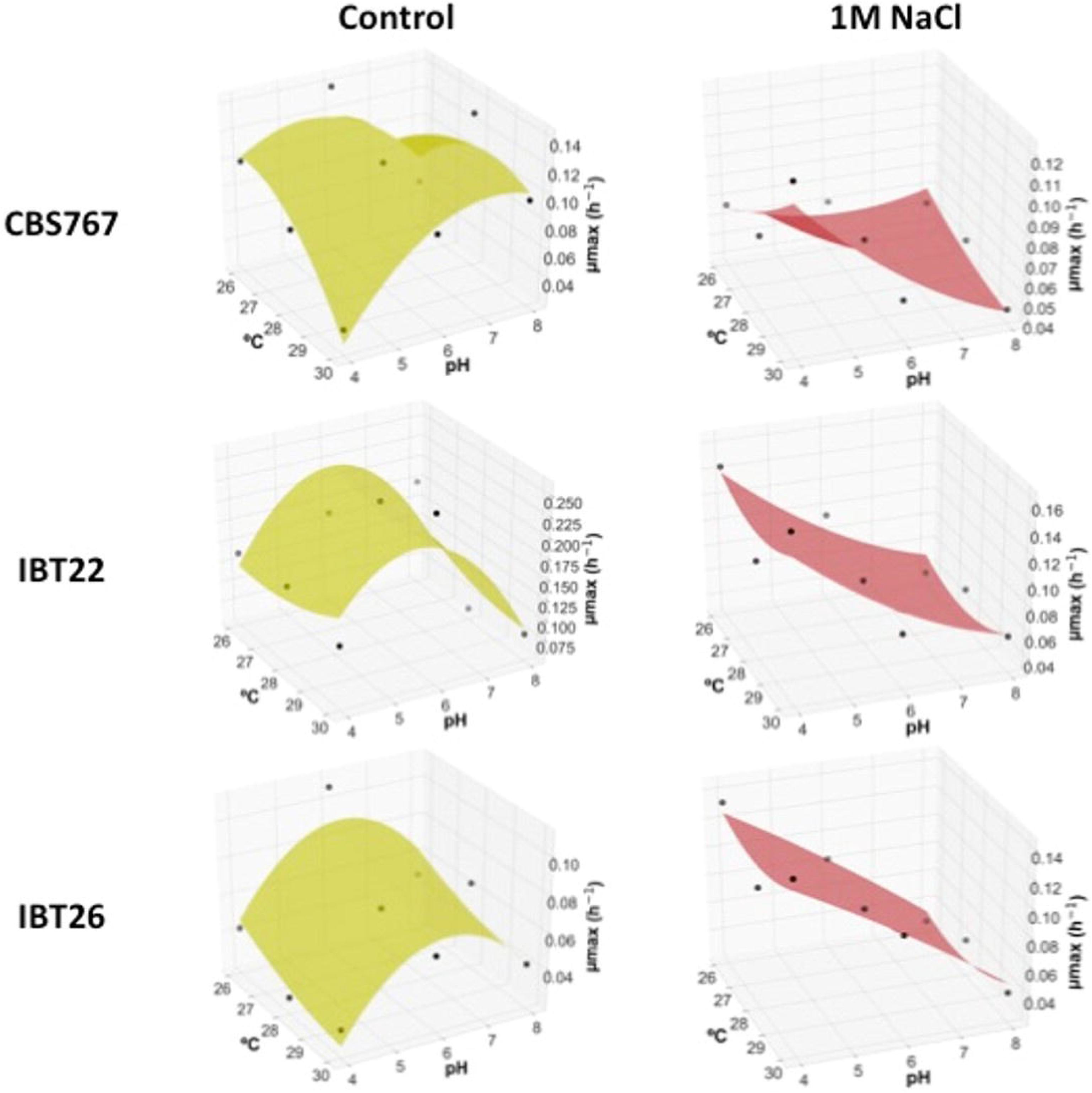
Maximum specific growth rates (μmax h^−1^) from micro-fermentations of *D. hansenii* strains CBS767, IBT22, and IBT26 in YNB control medium and supplemented with 1M NaCl (both with 2% glucose) as a function of pH and temperature (°C). The surface-response plots were generated using R.

As a result, we can observe that although the three *Debaryomyces* strains tested show a differential behaviour without the presence of salt in the culture media, once 1M of salt is present we can clearly observe how the cell growth at the different conditions shows a strikingly similar tendency for all the three strains (Figure 6). One important observation is that, as hypothesized by the results obtained from the 1L bioreactors, there is a positive correlation between high Na^+^ and low pH in terms of cell growth. This finally confirms what was already discussed in the previous section of this study and previously suggested by Almagro *et al.* (2000), about the summative effect of low pH and high salt. But the most important fact is that all these observations are extensive for the different *Debaryomyces* strains tested, hence confirming that the positive effect of high sodium is not strain dependent, but it can be inferred for *D. hansenii* in general.

## Discussion

The results obtained during this study, indicate that high salt concentrations in the culture media are indeed needed for optimal levels of cell performance in *Debaryomyces hansenii*. Especially in the case of NaCl, our results seem to finally confirm the halophilic behavior of this particular yeast: sodium does not exert detrimental effects over *D. hansenii*, but on the contrary. Another interesting finding is that NaCl exhibits a more significant positive impact than KCl, already evidenced in the spot-tests in solid media plates, but especially clear as the optimal growth rates obtained from the bioreactor experiments are reached in the presence of 1M of NaCl, closely followed by the values reached at 1M of KCl. This effect is even higher when the glucose source is scarce, as shown in the limited carbon (0.2%) bioreactor experiments.

There is, however, a longer lag phase when the yeast are grown in the presence of 2M NaCl, both seen in normal (2%) and limited (0.2%) carbon, which may have led to the previous conclusion by several other studies, performed in non-controlled environments, that the cell growth is affected by high sodium concentrations (Capusoni *et al.*, 2019). Although these previous conclusions can of course be due to the fact that those studies have been performed using different media and another yeast strain different than ours, and in non-controlled conditions (we actually see in our study that 2M salts in non-controlled conditions might point to a, mistakenly assessed, growth deficiency as observed in Fig. 4B and 4C), our data additionally show that the decrease in growth rate is not that dramatic at high salts (2M), and moreover, we also observed a slightly higher biomass yield in normal carbon, a much higher biomass yields in nutrient-limiting conditions compared to control conditions, as shown in Figure 2B, that the previous studies did not observe. This observation was again confirmed by spot-test studies with increasing salinity in solid media plates, in which we clearly observe that the presence of up to 1M salt (and particularly sodium) results in a faster growth compared with control conditions (no salt). Moreover, the presence of 2M sodium causes a slower growth compared to control, however after 240 hours the yeast has grown to the exact same levels than the control, hence there is no detrimental effect caused by high sodium concentrations: they simply grow slower and there is no inhibition of growth. Considering all of the above, we propose this as a survival strategy of *D. hansenii* to prevail in drying environments (e.g. during periods of drought). While other microbial species chose, as surviving strategy, to enter a dormant state or sporulate while environmental salinity levels are increasing (as a consequence of the lack of water), *Debaryomyces* reacts by increasing μ_max_ at moderate-high salinity levels (over 1M is already considered toxic for other yeast and bacterial species, so they have difficulties to proliferate (Yan et al., 2015)), hence increasing the glucose uptake rate (as our data reveals) thus ensuring getting the most of the available carbon for its survival in detriment to its competitors, and later when the salt concentration in the media keeps on increasing, slowing down the growth rate while still proliferating to overpopulate the area, changing the metabolic strategy from growing as fast as possible to growing as much as possible. This suggested strategy would be in accordance with the latest publications indicating that *Debaryomyces* are the most represented species in thawing Arctic glacier samples and coastal environments (Butinar *et al.* 2011; Jaques *et al.* 2014).

The dissolved oxygen levels in the culture medium decrease faster in presence of higher salt concentration, which suggests a higher metabolic activity. RQ values throughout the whole cultivation period remain below or equal at a value of 1 (reached constantly during the exponential phase), clearly supporting that there is no fermentative process, as stated previously. Finally, no ethanol is observed neither in the exometabolite analysis by HPLC, nor in the analysis in the off-gas by MS, therefore *D. hansenii* is herewith confirmed a crabtree negative yeast.

Our data without pH control in the bioreactor vessels indicate additional limiting factors during the cultivation, based on the CO_2_ production profiles, compared with pH controlled cultivations, evidenced by a long plateau phase, proving that shake flask experiments with non-controlled environment are not the ideal setup to obtain accurate conclusions about physiological and/or metabolic parameters, therefore we also suggest that previous studies providing conclusions obtained by this means, must be taken cautiously. It is also worth mentioning that it is not appropriate to choose randomly Na^+^ salts or K^+^ salts in order to study osmotolerance or halotolerance in general, as our results show that there is a clear differential effect exerted by either NaCl or KCl, as already suggested by Martinez *et al.* (2011 and 2012).

A very recent study, conducted by Ramos-Moreno *et al.* (2019) almost in parallel to ours, interestingly hypothesizes about a link between salt stress and oxidative stress response in *D. hansenii.* They cultured the yeast in the presence of high salt concentrations, reporting an increase in the expression of ROS detoxification genes (sodium showing a higher effect than potassium). Our findings align very well with their experimental results, and can complement their observations adding additional value: our study shows that the presence of salts (specially sodium) lead to a higher oxygen utilization. A higher respirative metabolism causes the activation of oxidative stress responsive genes, which would confirm the link between salt stress and the oxidative stress that Ramos-Moreno *et al.* suggest. In addition, both Ramos-Moreno *et al.* and our results confirm earlier studies pointing to a protective effect of NaCl against oxidative aggressions, such as the presence of H_2_O_2_ (Navarrete *et al.* 2009). In that case, the presence of NaCl would activate oxidative stress defenses as described above, thus they would be already active and therefore protective against the further aggression caused by addition of oxygen peroxide.

All in all, our results shed light upon the behaviour of *D. hansenii* in controlled bioreactor conditions, presenting this peculiar non-conventional yeast as a strain which is able to perform very well in “standard” cultivation conditions (no stress added) but whose performance gets significantly improved when environmental conditions get harsh: high salinity / osmotic pressure, media acidification and nutrient scarcity, all in combination. This undoubtedly confers to *D. hansenii* an incredibly strong potential for industrial production setups: those are the conditions that, upon large scale bioproduction processes (meaning vessels of 1000 litre or above) the microbial cells will encounter throughout the cultivation process, and that are limiting the suitable performance of microbes in bioreactors (Takors, R. 2012). Therefore, having such a strain with the above mentioned behaviour, and more importantly, obtaining sufficient information for understanding it (and, consequently, being able to take advantage of that knowledge) is of paramount interest for advancing in the field of cell factory design for industrial bioprocesses, and therefore for the biotech industry overall.

One of the strongest outcomes of our current study is the possibility of using *D. hansenii* in culture media containing relatively high salt concentrations, for industrial bioprocesses. On one hand, there is no need of using pure water sources, which significantly decreases the production costs as one could take advantage of desalination effluents, or even use directly sea water for the media composition, while still increasing the production yields (salinity will not affect *D. hansenii*, but will improve its cell performance, as shown by our research). On the other hand, using saline environments has another strong advantage which is the reduced risk of contaminations, hence sterilization costs would also be significantly reduced.

## Conclusion

Altogether, our findings reveal the beneficial role of salts, and more particularly sodium, in the cell performance of *Debaryomyces*, and open the need to further investigate how sodium and potassium influence the cell metabolism at a molecular level. It is clear that salts are not just tolerated in *D. hansenii*, but they play a crucial role in its survival strategy, to date underestimated. The presence of salts is needed for optimal cell performance. Further research, including a global expression analysis by RNAseq in steady-state continuous bioreactor cultivations, will shed light upon what are the intracellular mechanisms that trigger such metabolic changes, and how the discrimination between sodium and potassium occurs to trigger the different behavioural patterns described within this study. It will also be interesting from a biotechnological point of view, the identification of molecular elements that could potentially be responsive to the presence of salts, as our observations suggest that there are undoubtedly molecular switches which react to the presence or absence of sodium and/or potassium in the environment, triggering a specific metabolic response.

## Acknowledgements

We acknowledge the Novo Nordisk Fonden, within the framework of the Fermentation Based Biomanufacturing Initiative, for supporting this work. LRM received the fellowship “Ayuda de Movilidad Internacional para el Fomento de Tesis con Mención Internacional” from the University of Córdoba (Spain), to carry out her research work at DTU Bioengineering for three months. Prof. José Ramos (University of Córdoba, Spain) and Dr. Markus Bisschops (TU Delft, The Netherlands) are deeply acknowledged for critical reading of this manuscript, as well as Assoc. Prof. Christopher Workman for his assistance with the micro-fermentation data analysis. The authors would also like to thank the Fermentation Core at DTU Bioengineering, and Tina Johansen as well as Martin Nielsen for their technical support. This manuscript has been released as a pre-print at bioRxiv, (Navarrete *et al*.).

## Author’s contributions

JLM conceived the project. CN, ATF, LRM and MRK designed and performed the experiments. CN, ATF and MRK analyzed the data. CN and JLM wrote and revised the manuscript. All authors read, commented and approved the manuscript.

