## Supplementary figures for "A Physiological Characterization in Controlled Bioreactors Reveals a Novel Survival Strategy for *Debaryomyces hansenii* at High Salinity and Confirms its Halophilic Behavior": Supplementary material - Navarrete et al 2020.pptx

### Slide 1
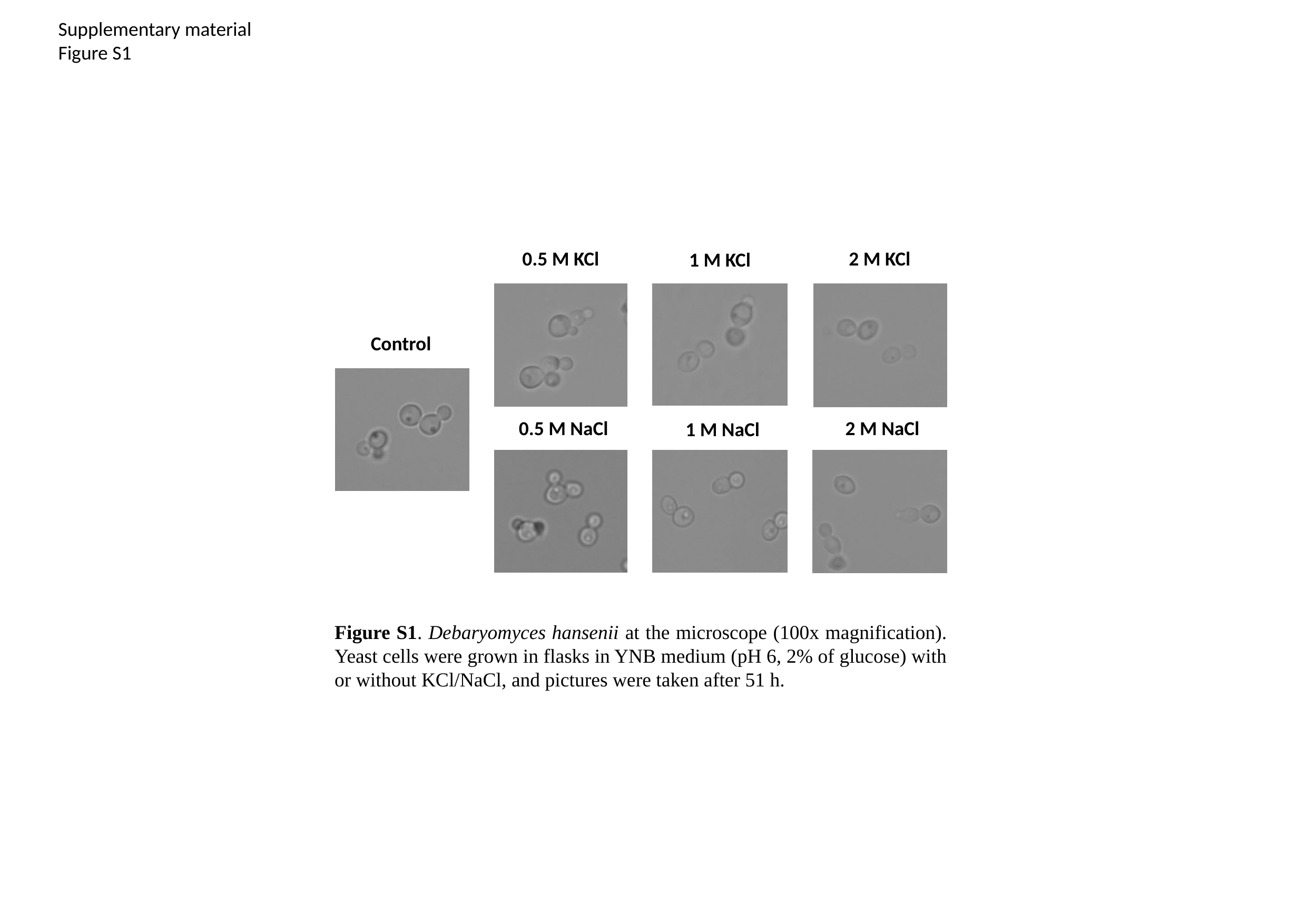

Supplementary material
Figure S1
0.5 M KCl
2 M KCl
1 M KCl
Control
0.5 M NaCl
2 M NaCl
1 M NaCl
Figure S1. Debaryomyces hansenii at the microscope (100x magnification). Yeast cells were grown in flasks in YNB medium (pH 6, 2% of glucose) with or without KCl/NaCl, and pictures were taken after 51 h.

### Slide 2
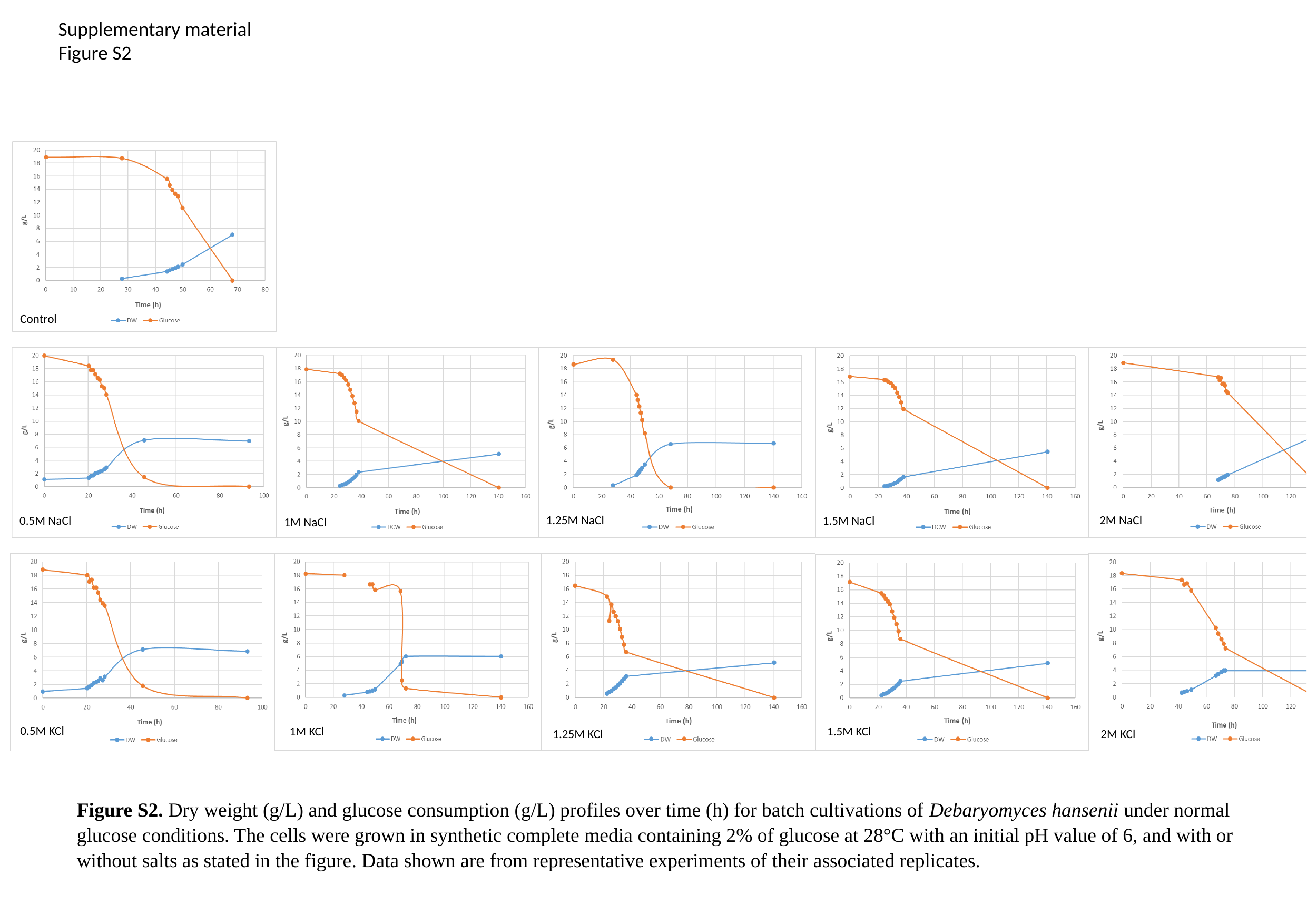

Supplementary material
Figure S2
Control
0.5M NaCl
1M NaCl
1.25M NaCl
2M NaCl
1.5M NaCl
0.5M KCl
1M KCl
1.25M KCl
2M KCl
1.5M KCl
Figure S2. Dry weight (g/L) and glucose consumption (g/L) profiles over time (h) for batch cultivations of Debaryomyces hansenii under normal glucose conditions. The cells were grown in synthetic complete media containing 2% of glucose at 28°C with an initial pH value of 6, and with or without salts as stated in the figure. Data shown are from representative experiments of their associated replicates.

### Slide 3
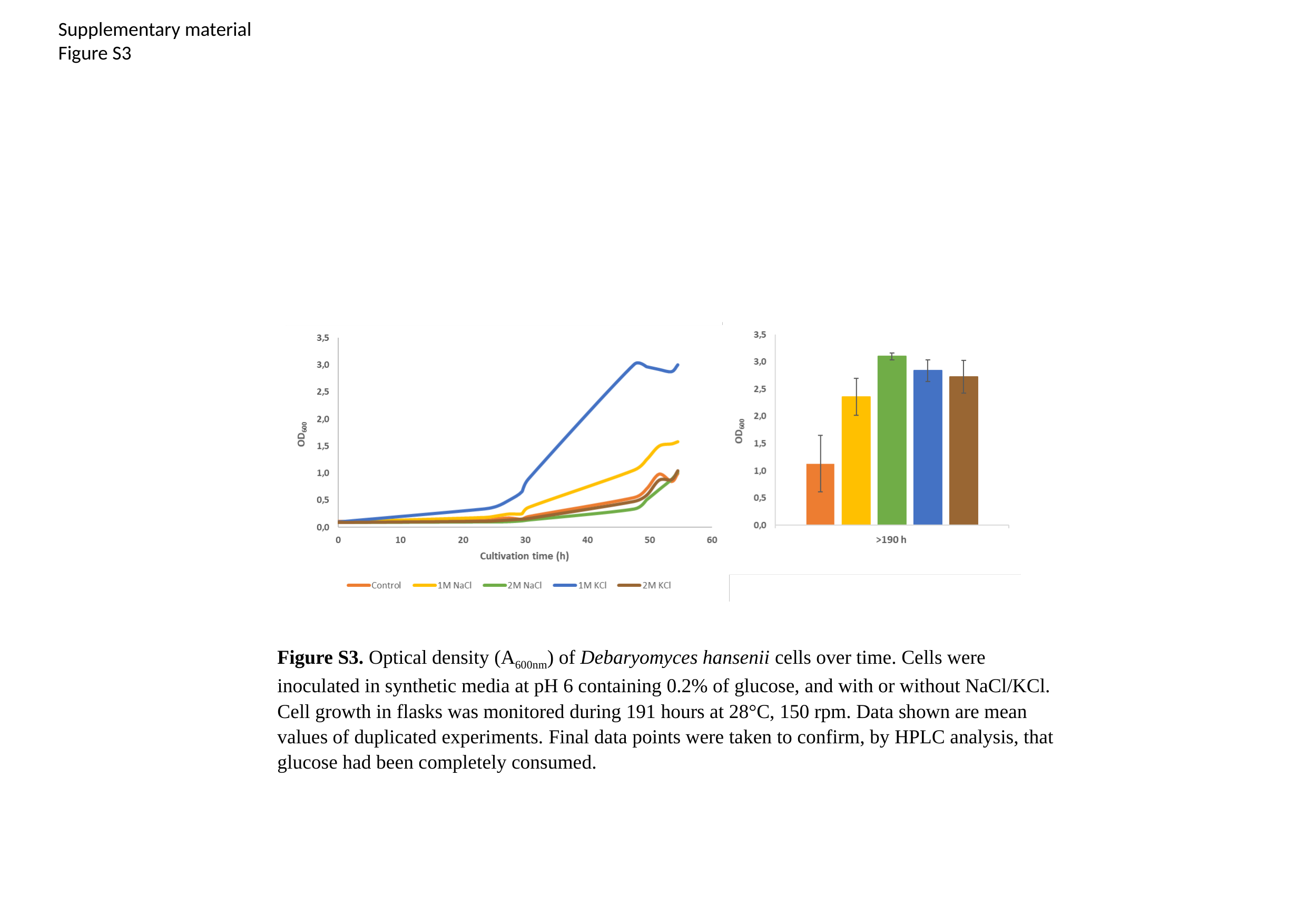

Supplementary material
Figure S3
Figure S3. Optical density (A_600nm_) of Debaryomyces hansenii cells over time. Cells were inoculated in synthetic media at pH 6 containing 0.2% of glucose, and with or without NaCl/KCl. Cell growth in flasks was monitored during 191 hours at 28°C, 150 rpm. Data shown are mean values of duplicated experiments. Final data points were taken to confirm, by HPLC analysis, that glucose had been completely consumed.

### Slide 4
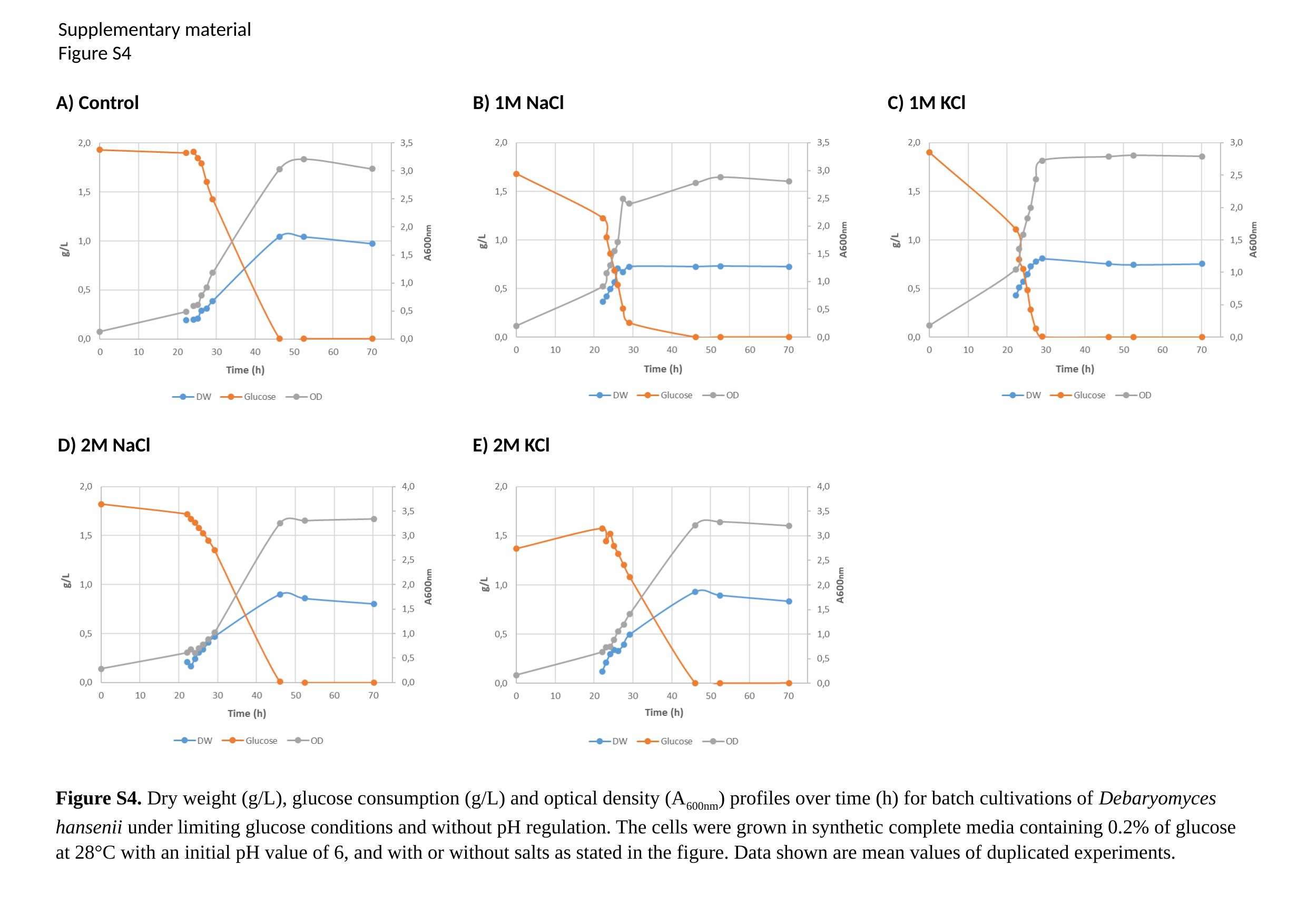

Supplementary material
Figure S4
C) 1M KCl
A) Control
B) 1M NaCl
D) 2M NaCl
E) 2M KCl
Figure S4. Dry weight (g/L), glucose consumption (g/L) and optical density (A_600nm_) profiles over time (h) for batch cultivations of Debaryomyces hansenii under limiting glucose conditions and without pH regulation. The cells were grown in synthetic complete media containing 0.2% of glucose at 28°C with an initial pH value of 6, and with or without salts as stated in the figure. Data shown are mean values of duplicated experiments.

### Slide 5
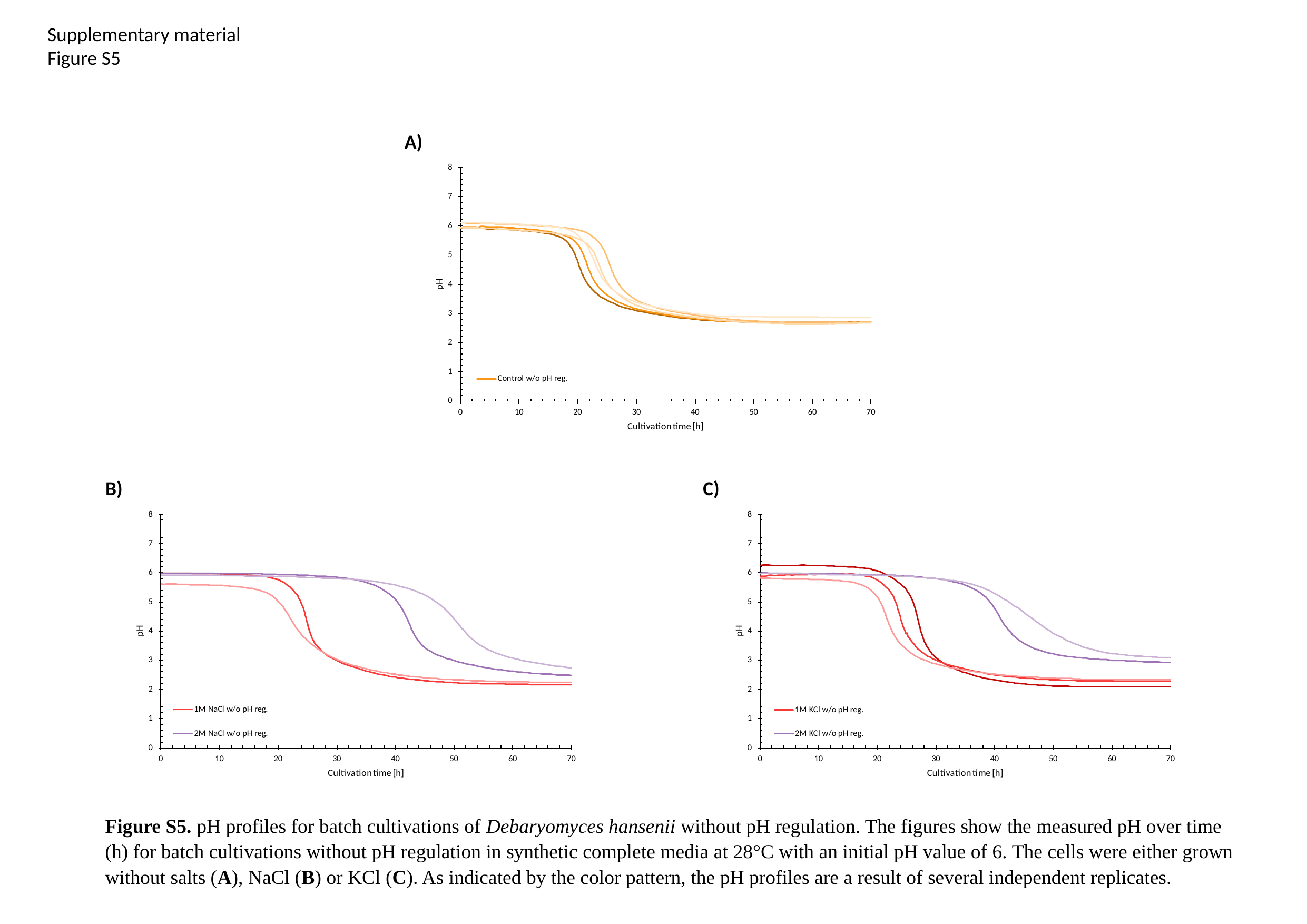

Supplementary material
Figure S5
A)
B)
C)
Figure S5. pH profiles for batch cultivations of Debaryomyces hansenii without pH regulation. The figures show the measured pH over time (h) for batch cultivations without pH regulation in synthetic complete media at 28°C with an initial pH value of 6. The cells were either grown without salts (A), NaCl (B) or KCl (C). As indicated by the color pattern, the pH profiles are a result of several independent replicates.

### Slide 6
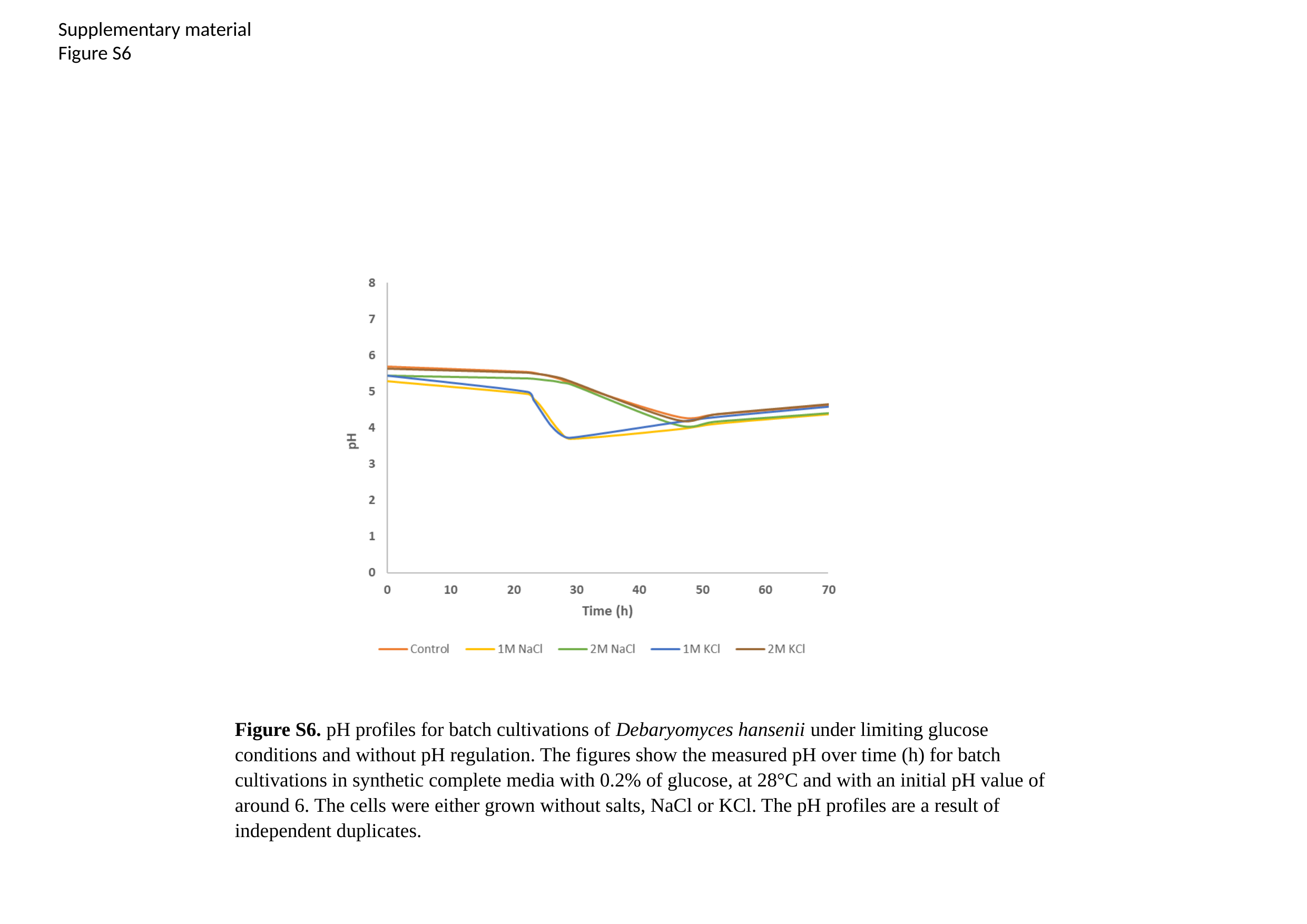

Supplementary material
Figure S6
Figure S6. pH profiles for batch cultivations of Debaryomyces hansenii under limiting glucose conditions and without pH regulation. The figures show the measured pH over time (h) for batch cultivations in synthetic complete media with 0.2% of glucose, at 28°C and with an initial pH value of around 6. The cells were either grown without salts, NaCl or KCl. The pH profiles are a result of independent duplicates.

### Slide 7
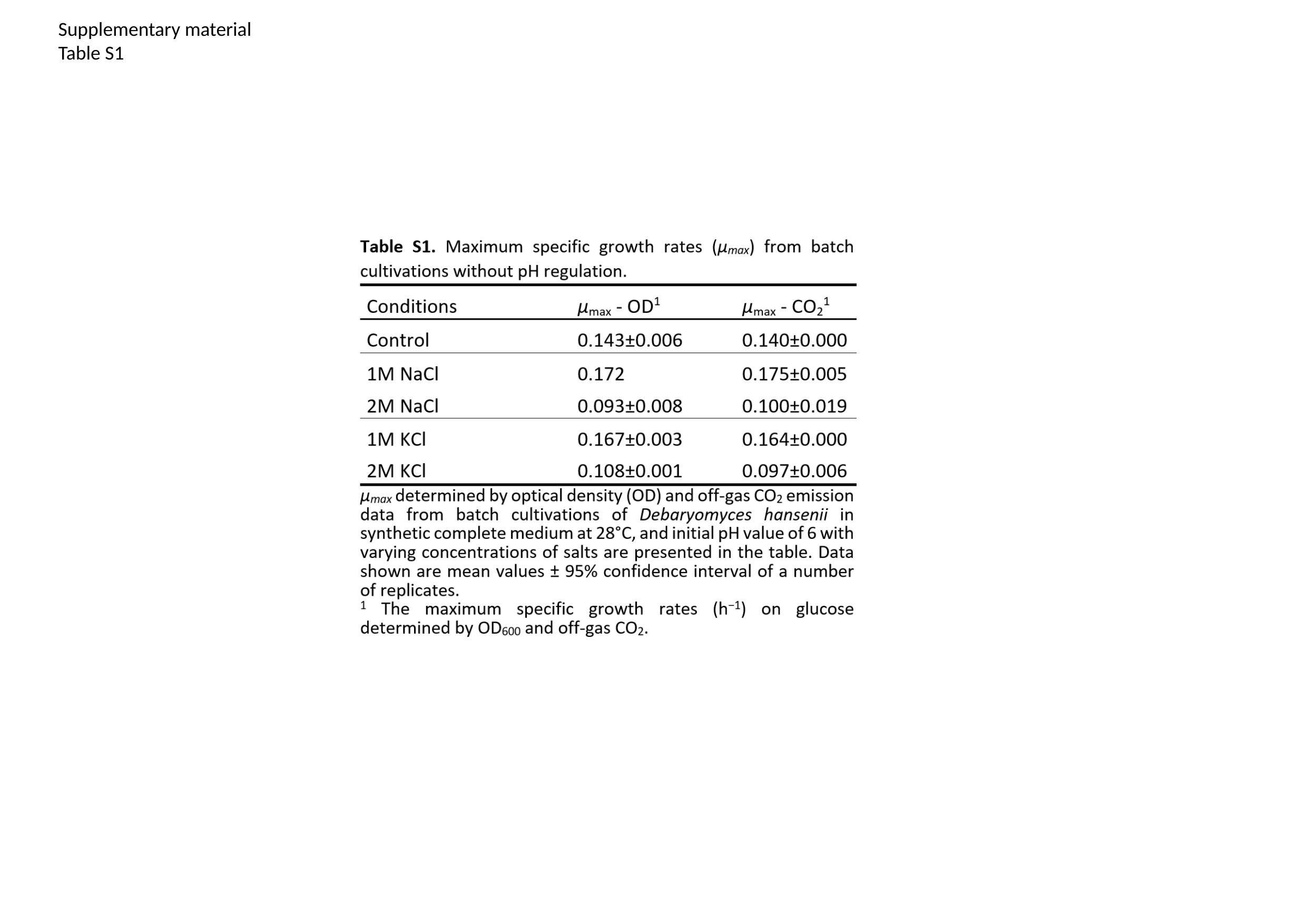

Supplementary material
Table S1
